# S3norm: simultaneous normalization of sequencing depth and signal-to-noise ratio in epigenomic data

**DOI:** 10.1101/506634

**Authors:** Guanjue Xiang, Cheryl A. Keller, Belinda Giardine, Lin An, Qunhua Li, Yu Zhang, Ross C. Hardison

**Author notes:** Corresponding authors: Yu Zhang, Department of Statistics, The Pennsylvania State University, Wartik Laboratory, University Park, Pennsylvania, 16802, USA,; Ross Hardison, Department of Biochemistry and Molecular Biology, The Pennsylvania State University, 304 Wartik Laboratory, University Park, Pennsylvania, 16802, USA.

## Abstract

Quantitative comparison of epigenomic data across multiple cell types or experimental conditions is a promising way to understand the biological functions of epigenetic modifications. However, differences in sequencing depth and signal-to-noise ratios in the data from different experiments can hinder our ability to identify real biological variation from raw epigenomic data. Proper normalization is required prior to data analysis to gain meaningful insights. Most existing methods for data normalization standardize signals by rescaling either background regions or peak regions, assuming that the same scale factor is applicable to both background and peak regions. While such methods adjust for differences in sequencing depths, they do not address differences in the signal-to-noise ratios across different experiments. We developed a new data normalization method, called S3norm, that normalizes the sequencing depths and signal-to-noise ratios across different data sets simultaneously by a monotonic nonlinear transformation. We show empirically that the epigenomic data normalized by our method, compared to existing methods, can better capture real biological variation, such as impact on gene expression regulation.

## INTRODUCTION

Epigenetic features of chromatin, such as histone modifications, transcription factor binding, and nuclease accessibility, play important roles in the regulation of gene expression. Advances in biochemical enrichment strategies and high-throughput sequencing technologies have made it possible to determine the landscape of epigenetic features at a genome-wide scale. In recent years, a large collection of genome-wide epigenetic profiles have been acquired in many cell types under different biological contexts (1–4). Quantitative comparison of these epigenetic profiles across different cell types is a powerful approach to study the biological functions of epigenetic modifications and infer functional elements in genomes. Technical heterogeneity across the data sets, such as differences in sequencing depth (SD) and signal-to-noise ratio (SNR), however, can create systematic biases that mask real biological variation (5). Proper data normalization is needed to correct these biases before meaningful insights can be gleaned from the data analyses (6, 7).

A commonly used strategy for data normalization is to calculate a scale factor between two data sets (8, 9), for example between a reference data set and a target data set, and then rescale the target data set according to the scale factor. The simplest scale factor is the ratio of the total signals between the two data sets, which we will refer to as TSnorm hereafter (Figure 1A). This approach is based on the assumption that the signals of a data set is dominated by the background regions and works well when real signals are scarce and take up only a small proportion of reads among the total. For epigenetic profiles, however, signal regions are often abundant, with drastically different number of peaks and reads across different data sets (9–12), whereas the background regions are more uniform across data sets. Recognizing this issue, some recent data normalization methods, such as SES and NCIS (13, 14), took a two-step approach. They first identify the background regions, and then they calculate the scale factor only from the background regions. While these methods can adjust for the scale differences in background regions between data sets, they implicitly assume that the same scale factor can be applied to peak regions as well. In reality, however, the signal-to-noise ratios between data sets are often different, thus the scale factor for the peak regions should be different from that for the background regions.

**Figure 1.**
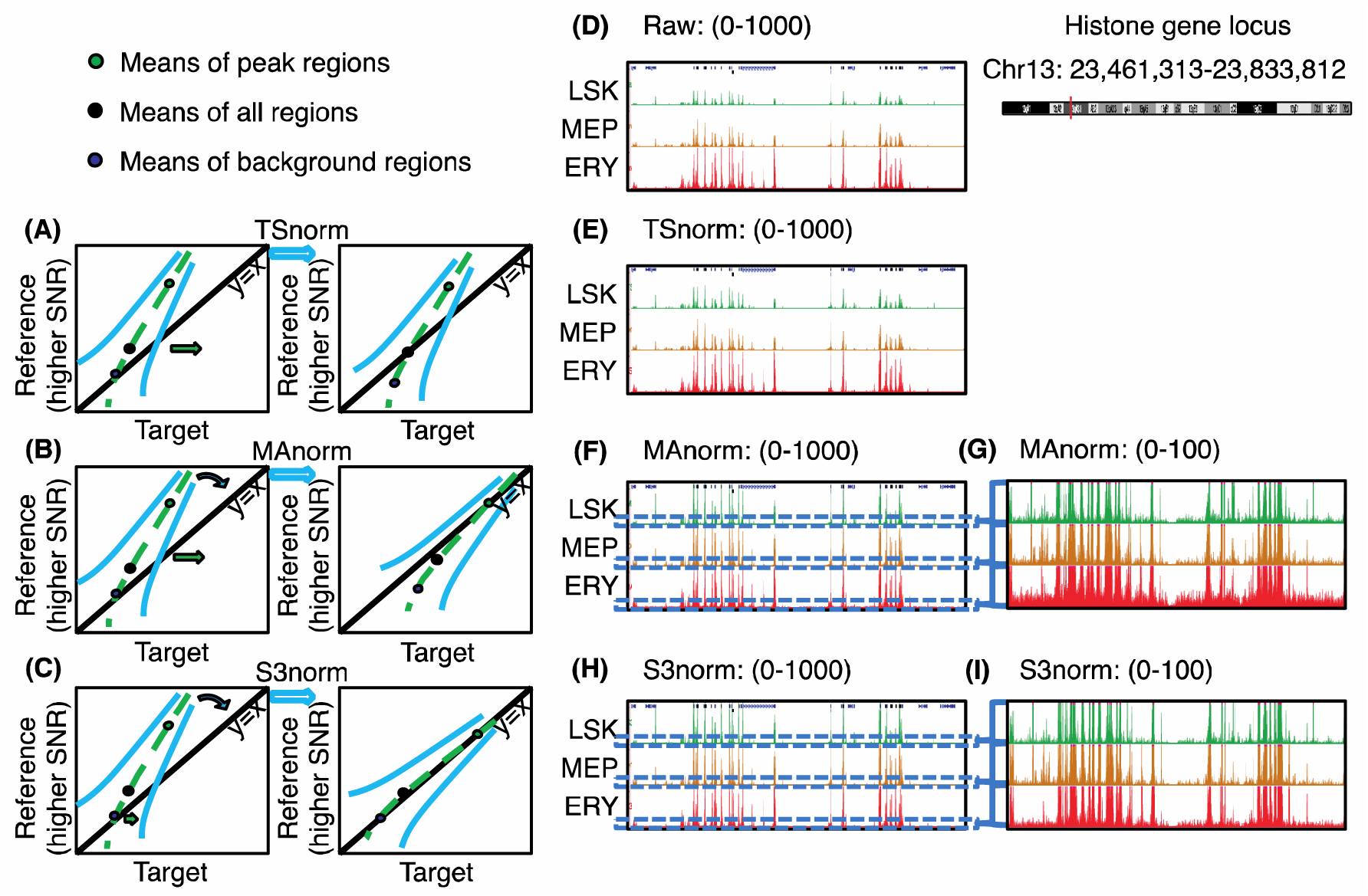
Impacts of different methods of data normalization. Panel **(A-C)**, respectively, shows schematic plots for the signals in two epigenomic data sets normalized by TSnorm, MAnorm, and S3norm. (**D-I**) The ATAC-seq or DNase-seq read counts at histone gene loci in three cell types (LSK, MEP, and ERY). Panel **(D)** shows the raw read counts in those three cell types. Panel **(E), (F)** and **(H)**, respectively, shows the read counts normalized by TSnorm, MAnorm, and S3norm. The scale of tracks is from 0 to 1000. Panel **(G)** and **(I)**, respectively, shows the zoomed-in version of the same regions in Panel **(F)** and **(H)**. The scale of tracks is from 0 to 100.

Some normalization methods focus on adjusting SNRs across data sets (15). MAnorm (6), one of the earliest methods to consider SNRs in ChIP-seq normalization, uses the MA plot (16) to fit a curve between signal intensity ratio (M) and average intensity (A) between data sets. The fitting is done using signals in the common peak regions between data sets, under the assumption that the normalized data sets in common peak regions should have the same SNRs. The fitted curve is then applied to adjust signals in peak regions (Figure 1B). MAnorm can adjust signals in peak regions, but not for background regions. It thus is not applicable for applications that utilize signals across the genome, such as genome segmentation (17–20). In segmentations, some epigenetic states are defined by low signals of features, in which case an increase of background noise could result in incorrect assignments to those states. The alternative methods LOWESS normalization and quantile normalization (QTnorm) have been used to adjust both SDs and SNRs by equalizing local signals between two data sets (21–23). When applying these two methods to data sets with substantially different numbers of peaks, they may increase background noise (or decrease peak signals) for data sets with fewer (or more) peaks (21). Finally, rank-based methods have been proposed to normalize data sets with different signal distributions by converting signals into ranks (24). Because they ignore the quantitative spread among signals, they may lose power, and therefore they are not considered in this study.

To illustrate the aforementioned issues encountered by the existing methods (Figure 1), we applied TSnorm and MAnorm to the nuclease accessibility data (ATAC-seq) at a histone gene locus in three hematopoietic cell types, namely, a stem and progenitor cell population (the lineage negative, Sca1 positive, c-kit negative cells or LSK), the megakaryocyte erythroid progenitor cells (MEP), and erythroblasts (ERY) (25). We chose this locus because active production of histones is required for cell replication, and histone genes usually have similar activities across all proliferating cell types. Thus, the profiles of nuclease accessibility in the neighborhood are expected to be similar across cell types, but the raw ATAC-seq signals in this locus were clearly weaker in LSK and MEP than in ERY (Figure 1D). After applying TSnorm, which used a single scale factor, the signals in LSK and MEP were increased but the signals of the peak regions in LSK and MEP were still weaker than in ERY (Figure 1E). This result is expected for a method that cannot match signals in both peak regions and background regions simultaneously. After applying MAnorm, which only used information in common peak regions to estimate a normalization model, the signals of the peak regions in both LSK and MEP were increased to match the level in ERY (Figure 1F), but the background was inflated in LSK and ERY (Figure 1G). These results illustrate the need for simultaneous adjustment of both peaks and background.

We developed a new two-factor normalization method, called S3norm, to <u>S</u>imultaneously normalize the <u>S</u>ignal in peak regions and the <u>S</u>ignal in background regions of epigenomic data sets. Unlike TSnorm or MAnorm, in which either background regions or common peak regions contribute to normalization, our method matches *both* the mean signals in the common peak regions *and* the non-zero mean signals in the common background regions between two data (Figure 1C), balancing the contribution of common peak regions and common background regions to data normalization. It matched the peak signals in our example data sets (ERY, LSK and MEP) without increasing noise the background signals (Figure 1H and I). In this paper, we demonstrate the superior performance of S3norm over existing methods using several epigenomic data sets with a wide range of data quality.

## MATERIALS AND METHODS

### Data preprocessing and evaluation

We used the data sets compiled by the **V**al**I**dated and **S**ystematic integrat**ION** of epigenomic data project (**VISION**: usevision.org), which includes eight epigenetic marks (H3K4me3, H3K4me1, H3K27ac, H3K36me3, H3K27me3, H3K9me3, CTCF occupancy, and nuclease sensitivity) in twenty hematopoietic cell types of mice (20, 26–28). Using the bam files processed by the pipeline of VISION project as the input data (20, 28, 29), we divided the mm10 mouse genome assembly into ∼13 million 200-bp bins and counted the number of reads mapped to each bin (30). The reads counts per bin comprised the raw signals for each data set. The bin size of 200-bp was selected based on previous studies that considered this bin size as approximately the size of a nucleosome plus spacer (19, 31). For each data set, the SD was estimated by the number of mapped reads, and the SNR was estimated as the Fraction of Reads in Peaks (FRiP score) (32). The VISION data sets were generated in different laboratories at different times using different technologies, leading to substantial variation in signal quality across data sets (Supplementary Figure 1). Considering H3K4me3 experiments as an example, the total number of mapped reads ranged from <1 million to >10 millions, and the FRiP score ranged from <0.1 to >0.9. This large variability in both SDs and SNRs requires both aspects to be properly normalized to enable meaningful downstream analysis. Indeed, this large variation in both SD and SNR served as a motivating problem to develop our normalization method.

### Simultaneous normalization of both peak regions and background regions

S3norm is a normalization method that aims to simultaneously match the SD and the SNR between a target data set and a reference data set. This aim is achieved by applying a two-factor nonlinear transformation to match both the mean signal in the common peak regions and the non-zero mean signal in the common background regions between the two data sets (Figure 2). Because the numbers and the signals of the unique peaks can differ between the two data sets, S3norm learns its normalization parameters only from the signals in the common peak regions (6) and the signals in the common background regions. S3norm is built on two assumptions derived from the biological principles of epigenetic events. First, we assumed that epigenetic events shared by multiple cell types tend to regulate processes occurring in all those cell types, such as expression of constitutively active genes, so that the mean signal of common peaks should be the same after normalization. Second, we assumed that the signals in common background regions are technical noise, and thus, they should be equalized after normalization. Based on these two assumptions, S3norm matches the mean signals in the common peak regions and the non-zero mean signals in the common background regions between the two data sets. Here we used the non-zero mean instead of the mean signal because some epigenomic sequencing data sets have a large number of bins with values of zero (33). Inclusion of the large numbers of zeros generates low mean signals that are problematic outliers compared to other datasets that have fewer bins with values of zero. S3norm can also work for more than two data sets, in which case the common peak regions and the common background regions are those shared by all data sets.

**Figure 2.**
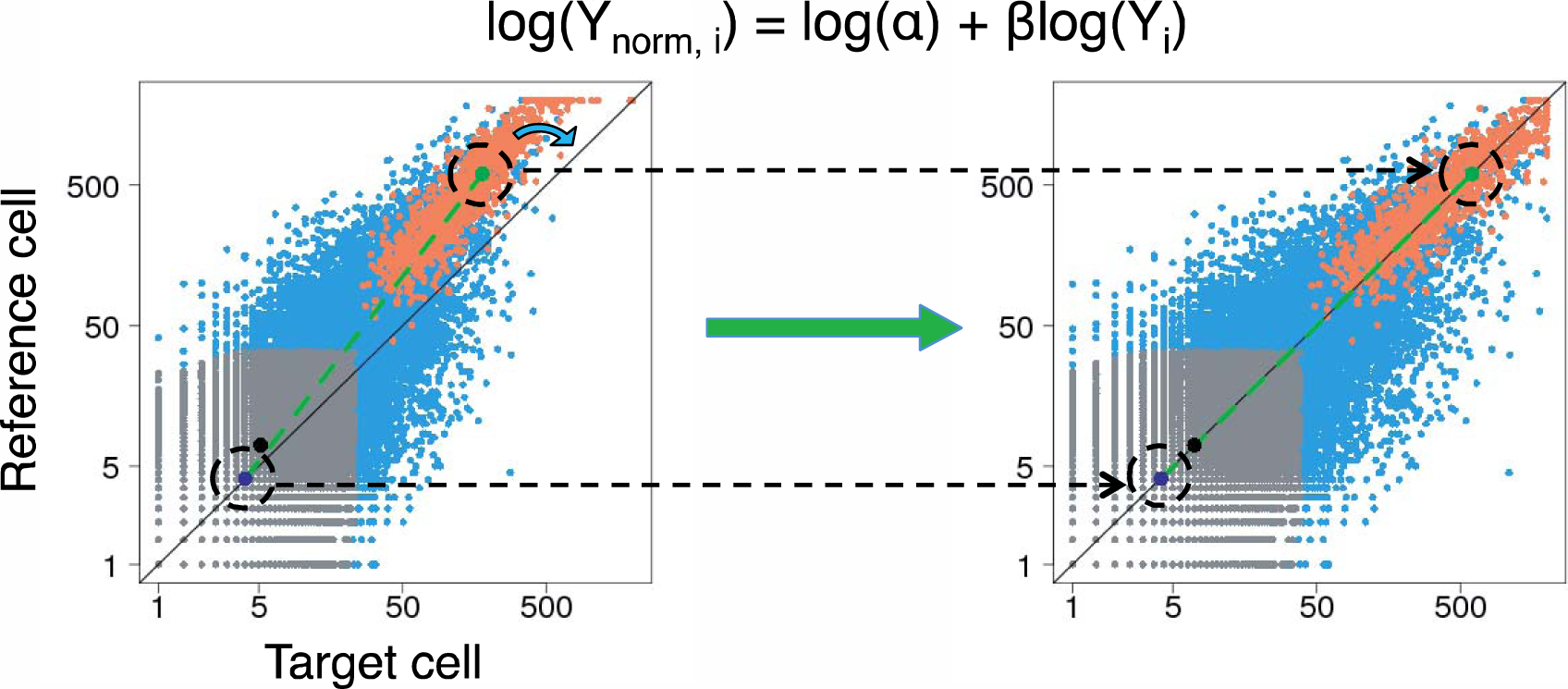
Overview of the S3norm method. The graphs present scatterplots of read counts (log scale) in 10,000 randomly selected genome locations (200bp) in target cell (x-axis) and reference cell (y-axis). The left figure is the signal before S3norm. The right figure is the signal after S3norm. The S3norm applies a monotonic nonlinear model (log (Y_norm,i_) = log(α) + βlog(Y_i_)) to rotate the target signal so that (1) the mean signals of common peaks (green point, highlighted by black dash circle) and (2) the mean signals of common background (dark blue point, highlighted by black dash circle) can be matched between the two data sets. The original data were split into three groups: the common peak regions (orange), the common background regions (gray), and the remaining bins (blue). The overall mean is represented by a black point.

To match the mean signals, we treat one data set as the reference and the other data set as the target, transforming the signals in the target data set by the following monotonic nonlinear function:

Let Y_i_ and Y_norm,i_ denote the signal of bin *i* in the target data set before and after normalization, then

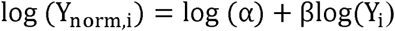

where α and β are two positive parameters to be learned from the data. Specifically, α is a scale factor that shifts the signals of the target data set in log scale, and β is a power transformation parameter that rotates the signals of the target data set in log scale (Figure 2). There is one and only one set of values for α and β that can simultaneously match the mean signals in the common peak regions *and* the non-zero mean signals in common background regions between the two data sets. Our approach solves the values for α and β which satisfy the following two equations, so that the mean signals in the common peak regions *and* the non-zero mean in the common background regions can be matched between two data sets:

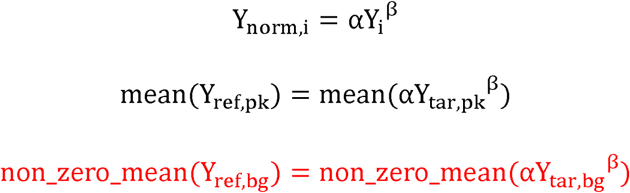

where the mean(αY_tar,pk_^β^) is the mean of normalized signals in common peak regions and the non_zero_mean(αY_tar,bg_^β^) is the non-zero mean of normalized signals in common background region, respectively, in the target data set. Likewise, the mean(Y_ref,pk_) is the mean of normalized signals in common peak regions and the non_zero_mean(αY_ref,bg_^β^) is the non-zero mean of normalized signals in common background region, respectively, in the reference data set. The values of α and β were estimated by the Newton-Raphson method (34).

For the reference data set, we choose the data set with the best SNR as the reference. In the S3norm package, users also have options to choose another data set *or* generate a reference data set by using the median (or mean) signal of all data sets for each genome position.

The common peak regions and the common background regions for S3norm to learn the non-linear transformation model were defined by using FDR = 0.1 as the threshold. As shown in Supplementary Figure 2A, the peaks shared by more cell types are more robust to the FDR threshold.

Using these common peak regions, we also tested our assumption that epigenetic events shared by multiple cell types tend to regulate processes occurring in all those cell types, and thus the epigenetic datasets should have similar mean signals. We found the genes associated (by proximity) with the ATAC-seq peaks shared by all 18 cell types in this analysis were significantly enriched in GO terms about nuclear processes common to all cells, such as chromatin assembly, DNA replication, and RNA metabolism (35). Thus, using these regions for S3norm to learn the non-linear transformation model fits well with our assumption.

### Generating signal tracks from normalized signal

To facilitate use in downstream analysis, such as peak calling and genome segmentation, we generated signal tracks (bigwig format) of the S3norm normalized signals. A script for generating these signal tracks is provided at GitHub. We followed a similar method as the one adopted in MACS (9) except that the Poisson model used to adjust for fluctuation in the local background (36–39) was replaced by a Negative Binomial (NB) model. In ChIP-seq data, the variance is often greater than the mean (supplementary Figure 3), so a NB is preferred as the background model because it estimates the mean and variance separately, whereas the Poisson model has the same mean and variance.

For ChIP-seq, there are usually two data sets for each experiment, one is referred as an immunoprecipitation (IP) sample, which is a data set generated by sequencing the DNA after immunoprecipitation by target-specific antibody, and the other one is the corresponding control sample, which is another data set generated by sequencing either the input DNA without immunoprecipitation or the DNA after immunoprecipitation by non-specific antibody.

The NB background model was defined as follows. Let r_i_ and 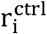 denote the read counts in bin i in a IP and a control, respectively. Let M and σ^2^ denote the mean and variance of read counts in the IP in the common background regions, and M^ctrl^ denote the mean read counts in the control in the common background regions. The dynamic NB background model is defined as:

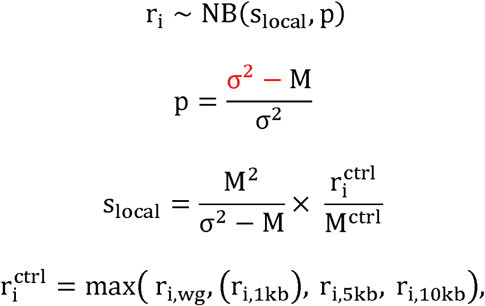

where p denotes the probability of success parameter in the NB model, and s_local_ denotes a shape parameter of the NB model. For each bin i, s_local_ is adjusted by 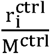 to capture any local bias as reflected in the control. The increase of control signal 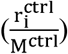 is equivalent to the decrease of σ^2^ which can generate a more significant p-value. The 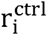 is the local mean read count learned from the control computed in the same way as in MACS (9). The r_i,wg_ is the genome mean read count in the control, the r_i,1kb_, r_i,Skb_ and r_i,10kb_, are mean read counts of different window sizes centered at the bin i in the control. The local mean read counts are calculated as the maximum of r_i,wg_, r_i,1kb_, r_i,Skb_ and r_i,10kb_. For data sets without a control (i.e. ATAC-seq), the value for 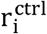 can be generated with both of two modifications (9). First, the 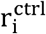 and M^ctrl^ are estimated from the IP instead of the control. Second, the r_i,1kb_ is not used to estimate 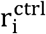.

Just like we avoided bins with values of zero in the S3norm normalization (previous section), we also only used the non-zero data to estimate the p and the s_local_ in the dynamic NB model.

Let NM1 and NM2 denote the non-zero mean and non-zero mean of square of the read counts of all bins, respectively. The p and the s_local_ in the dynamic NB model can be defined as follows (see Supplementary Methods for details):

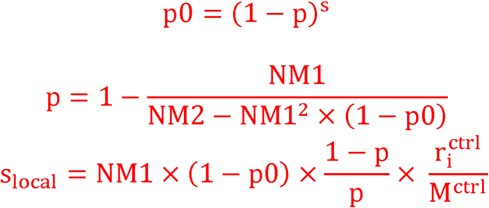

Finally, -log10 p-value of read count per bin, as derived from the NB background model, is used as the processed signal in the S3norm signal track.

### Predicting gene expression from histone modifications

Previous studies have shown that histone modifications such as H3K4me3 and H3K27ac are strong predictors of gene expression (40). We assumed properly normalized histone modifications data can accurately predict gene expression. Following the study design in Dong et al 2012 (41), we used ten-fold cross validation to evaluate the predictability of gene expression. For each cross validation, we randomly selected 90% of the genes as training genes and the remaining 10% of the genes as the testing genes. We first trained a regression model to predict expression of training genes in one cell type (training cell type). We then applied the trained model to predict the expression of testing genes in another cell type (testing cell type). The Reads Per Kilobase of transcript per Million mapped reads (RPKM) in log2 scale was used as the estimate of gene expression. The histone modification signal was defined as the mean read counts of the histone modification in a 5kb window centered at transcription start site (TSS). The predictability of gene expression was measured by mean square error (MSE) between the observed gene expression and the predicted gene expression in the testing genes in the testing cell type. To prevent a bias from a specific regression model, we performed this analysis by using four different commonly used regression models, specifically a local regression (loess) model, the 2-step linear regression model (41), a linear regression model, and a support vector machine regression (SVR) model.

### Calling peaks from epigenomic data by MACS2

To compare of the influence of data normalization on peak calling, we applied MACS2 to call peaks from CTCF ChIP-seq data normalized by different normalization methods. We first generated the signal tracks in each cell type. For all methods, the signal track was generated by the -log10 p-value (input signal for bdgpeakcall in MACS2 package) of normalized reads count based on the previously described NB background model. We then used the bdgpeakcall in the MACS2 package in the default setting to call peaks from the signal tracks. The threshold was FDR = 1e-2. For each normalization method, the CTCF peaks were first called in 11 cell types that have CTCF occupancy data in VISION project. We used the UpSet method (42) to visualize the number of commonly called peaks from different normalization methods.

To evaluate the type I error (false positive peaks) in these CTCF peak calling results, we compared both the proportion of peaks with a CTCF binding site motif (Jaspar id: MA0139.1) (43) and the signal consistency between the biological replicates in these peaks. For the proportion of peaks with CTCF binding site motif, we used FIMO in its default setting to scan for the CTCF binding site motif (Jaspar id: MA0139.1) (43) in those peaks.

The signal consistency between the biological replicates was measured by the mean square error (MSE) between the two biological replicates.

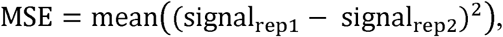

where signal_rep1_ and signal_rep2_ are the signals in two biological replicates. The false positive peaks tend to be the peaks with lower signal. To measure the signal consistency of peak with different signal levels, we calculated the cumulative MSEs.

### Generating ATAC-seq signal strength state by IDEAS

To compare the impact of normalization by QTnorm and S3norm (19) in a context in which the background signal had an influence, we measured the impact of each normalization on genome segmentation. Specifically, we used a simple application of the IDEAS genome segmentation algorithm to generate states defined by the ATAC-seq signal strength across the whole genome (19). We first normalized the raw read counts by the QTnorm and S3norm. We then computed the -log10 p-value based on the dynamic NB background model for each 200bp bin and used them as the input signal for IDEAS. The IDEAS algorithm learned the most commonly occurring states of ATAC-seq signal strength and assigned each 200bp bin along the genome into one of those signal strength states. For this analysis, we kept the biological replicates separate so that the consistency of the state assignments between the two biological replicates could be used to evaluate the performance of the results. We used the Adjusted Rand Index (ARIs) to measure the consistency of the state assignment between each pair of two biological replicates.

## RESULTS

### S3norm overview

We introduce a new data normalization method called S3norm that uses a nonlinear transformation to normalize signals in both peak regions and background regions simultaneously. The goal of the S3norm method is to match *both* the mean signal in the common peak regions *and* the non-zero mean signal in the common background regions between a target data set and a reference data set, which is the data set with the best SNR (Figure 2). The method employs a nonlinear transformation model with parameters learned from the signal in both common peaks and common background regions. This nonlinear transformation can rotate the target signal in log scale to achieve simultaneously the desired matches with the mean signals in both the common peak regions and common background regions of a reference data set. As a result, the method can boost signals in peak regions in the target data set without increasing the background noise, thereby increasing the SNR in the target data sets (see Materials and Methods for details). Because it does not inflate the background signal, S3norm is well suited as a normalization tool for genomic analyses that involve both peaks and background, such as segmentation of epigenetic states.

### Evaluation by ATAC-seq

We first compared the performance of S3norm and other normalization methods for their abilities to match the signal in both peak regions and background regions. We used the ATAC-seq data sets in immature megakaryocytic (iMK) cells (∼92 million reads for replicate 1 and ∼74 million reads for replicate 2) and LSK cells (∼53 million reads for replicate 1 and ∼59 million reads for replicate 2) because they have quite different SNRs (Figure 3A; the mean signal for common peaks in iMK was about 3.7-fold greater than that in LSK). We used the iMK dataset as the reference because it had a higher SNR. For all normalization methods, the signal of the target data set was matched to the signal of the reference data set.

Because the two data sets had similar signal in background regions, the TSnorm method had little impact on the results (Figure 3B), i.e., the peak signals in iMK remained consistently higher than those in LSK after TSnorm normalization. The MAnorm method did normalize the signals in peak regions to match peak signals in the LSK and iMK data sets (Figure 3C). However, MAnorm increased the signal in background regions in the LSK data set. The poor performance of TSnorm and MAnorm was expected, as they used either background regions or peak regions to calculate scale factors, but neither considered both types of regions simultaneously. In contrast, normalization by QTnorm and S3norm included signals in both peak regions and background regions and generated similar mean signals of the common peak regions (green point) and the common background regions (dark blue point) between the two data sets (Figure 3D and E).

**Figure 3.**
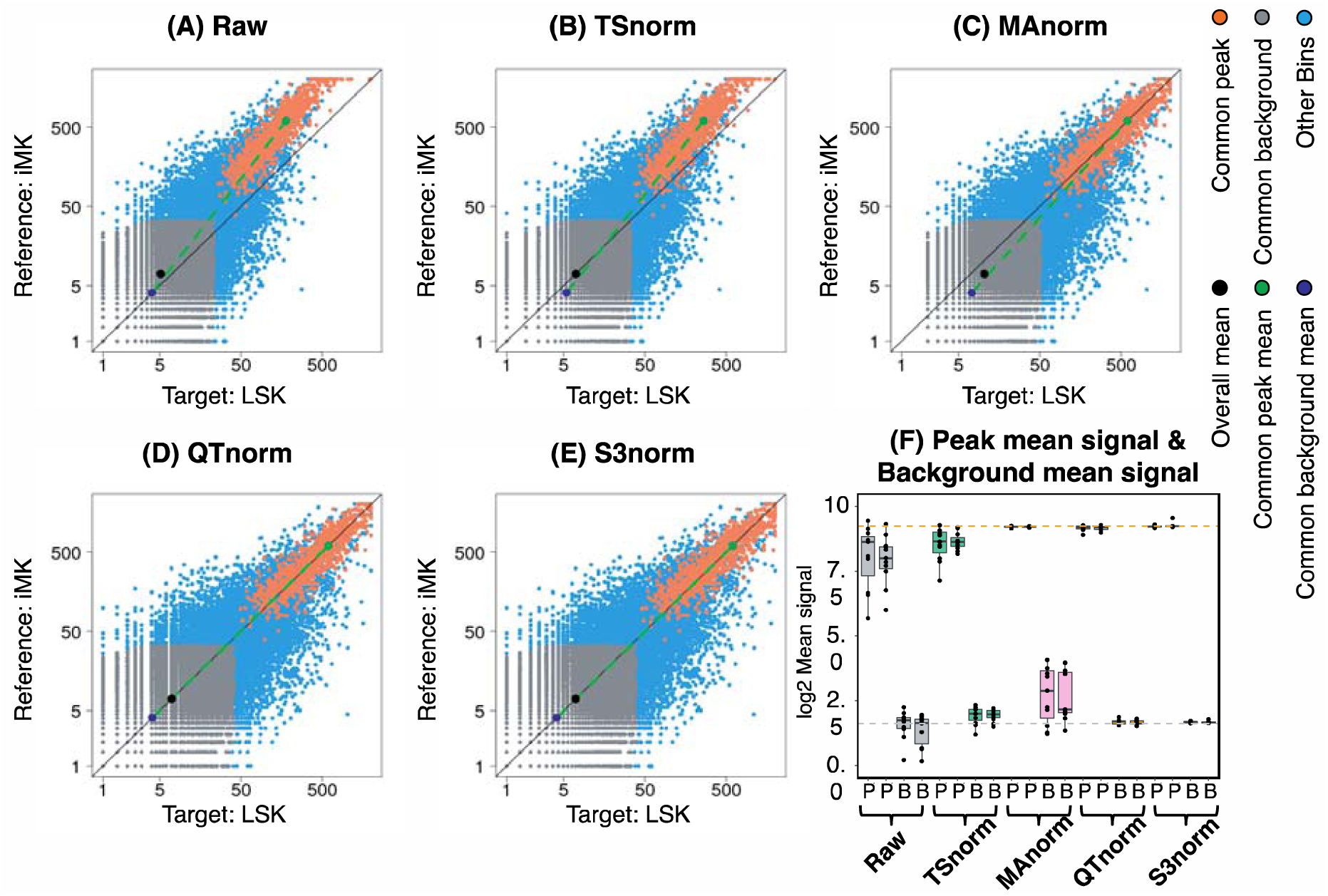
Comparison of normalization methods on peaks and background in ATAC-seq experiments. The scatterplots of ATAC-seq signal in iMK (reference data set) and LSK (target data set) are shown on a log scale. **(A)** The scatterplot of the raw signal between reference data set and target data set. Panel **(B)** to **(E)**, respectively, shows the scatterplot of the signal after TSnorm, MAnorm, QTnorm, and S3norm. **(F)** The boxplot of the mean signals in common peak regions (**P**) and the mean signals in the common background regions (**B**) in the biological replicates of different cell types.

We next compared the four normalization methods for their impact on the similarity of replicates, expecting that a more suitable normalization would generate closer matches between replicates. The four methods were used to normalize all ATAC-seq data sets in the VISION project that have biological replicates, employing the data set with the highest SNR (iMK) as the reference. The signals in the common peak regions showed substantial differences both between replicates and across all data sets after TSnorm (Figure 3F), illustrating the limitation of single factor normalization. MAnorm adjusted the signal in common peak regions appropriately, producing close similarities in distributions between replicates, but the signals in the background regions became more heterogeneous than they were before normalization. In contrast, S3norm and QTnorm effectively adjusted the signals in both types of regions so that the normalized signals became much more comparable both between replicates and across data sets.

### Evaluation by gene expression

When properly normalized, histone modification data should enable a more accurate prediction of gene expression than achieved with inadequately normalized data. Thus, we evaluated the effectiveness of different normalizations by analyzing the ability of histone modification data normalized by each method to accurately predict gene expression. Modeling approaches have been used to predict gene expression from histone modifications, and the quantitative relationships learned from one cell type can be applied to predict gene expression in other cell types (41, 44). The predictability, however, will be reduced if the epigenomic data across cell types are not properly normalized. We thus use the predictability of gene expression from different normalized epigenetic data to evaluate their ability to reflect real biological variation.

Specifically, we randomly selected 90% of genes (*Training Genes*) to train four commonly used regression models (loess regression, 2-step linear regression (41), linear regression, and SVR) to predict gene expression from H3K4me3 and H3K27ac normalized signals around a gene TSS. We then evaluated the performance of these regression models on the remaining 10% of genes (*Testing Genes*). We first trained the regression models using the *Training Genes* in one cell type (*Training Cell type*) and then evaluated the models using the *Testing Genes* in both *Training Cell type* and a different cell type (*Testing Cell type*).

The evaluation in the *Training Cell type* ascertained whether regression models can successfully learn robust quantitative relationships between gene expression and normalized signals for histone modifications. All normalization methods generated data that gave good performance for lowess regression (Figure 4A), shown by the very low mean squared errors (MSEs) in the *Training Cell types*. A similarly uniform good performance for all normalization methods was observed for three other regression models (Supplementary figure 4A-C).

**Figure 4.**
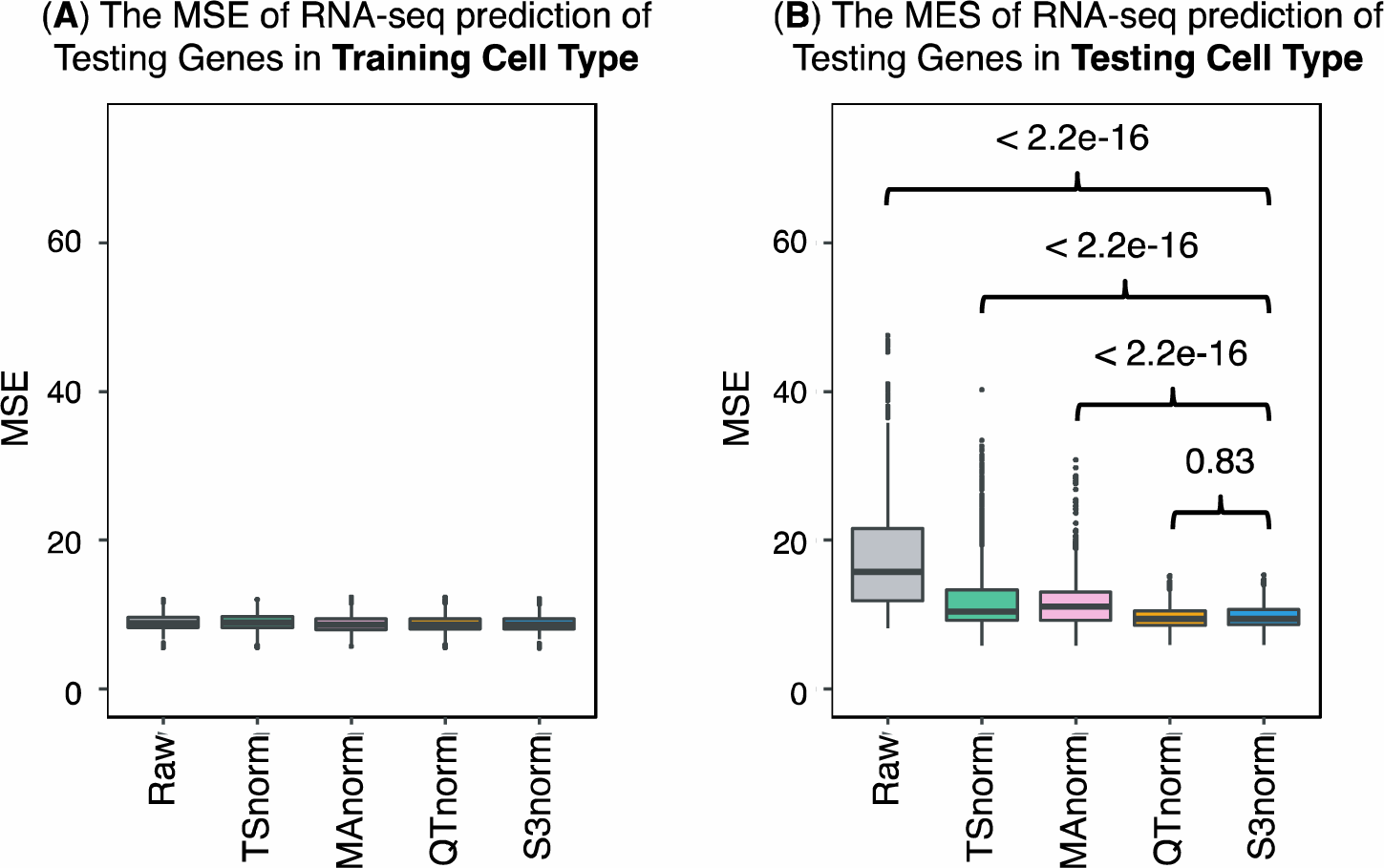
Comparing S3norm and other methods by their ability to predict gene expression. (**A**) The MSE of the observed RNA-seq signal and the predicted RNA-seq in ten-fold cross validation in the **Training Cell Type** by using a loess regression model. (**B**) The MSE of the observed RNA-seq signal and the predicted RNA-seq in ten-fold cross validation in the **Testing Cell Type**. The p-values above the boxes come from the Wilcoxon test that tests if the MSE of S3norm are significantly better than the other methods.

Then we evaluated whether the learned quantitative relationships can be applied to different cell types by comparing the performances of the trained models on the *Testing Genes* in the *Testing Cell Type*. As shown in figure 4B and Supplementary figure 4D-F, the models trained on S3norm signals and QTnorm signals always performed better (Wilcoxon test p-value < 1e-4) than the models trained on the TSnorm signals and MAnorm signals. This result shows that the quantitative relationships learned from the histone modification signals normalized by QTnorm and S3norm transferred well to other cell types, thus indicating a more biologically meaningful normalization result.

### Evaluation by peak calling and signal consistency

The previous evaluations showed the superior performance of S3norm and QTnorm than some other normalization methods. Both methods match the signal in peak regions and background regions simultaneously. However, QTnorm differs from S3norm in its assumption that normalized signals have the same distribution across data sets. This assumption is particularly questionable for epigenomic data, because the number of epigenetic peaks usually differs substantially across cell types. If two data sets have different numbers of peaks, QTnorm would force some background signals in the data set with fewer peaks to match the peak signals with the same rank in the data set with more peaks, potentially creating false positive peaks. Therefore, we evaluated the impact of each normalization method on identifying more meaningful peaks in ChIP-seq and ATAC-seq datasets.

In this evaluation, we first called peaks on CTCF ChIP-seq data from the VISION project using the signal normalized by different methods. We first compared the number of peaks overlapping between sets by using the UpSet method (Figure 5A, first panel) (42). As expected, a large number (78,789) of peaks were called consistently on the data normalized by all methods, but almost as many (67,444) peaks were called only from the QTnorm signal. These peak calls that were unique to normalization by QTnorm could be false positive peaks created by forcing identical distributions across data sets and thereby inflating the background such peaks are called erroneously.

**Figure 5.**
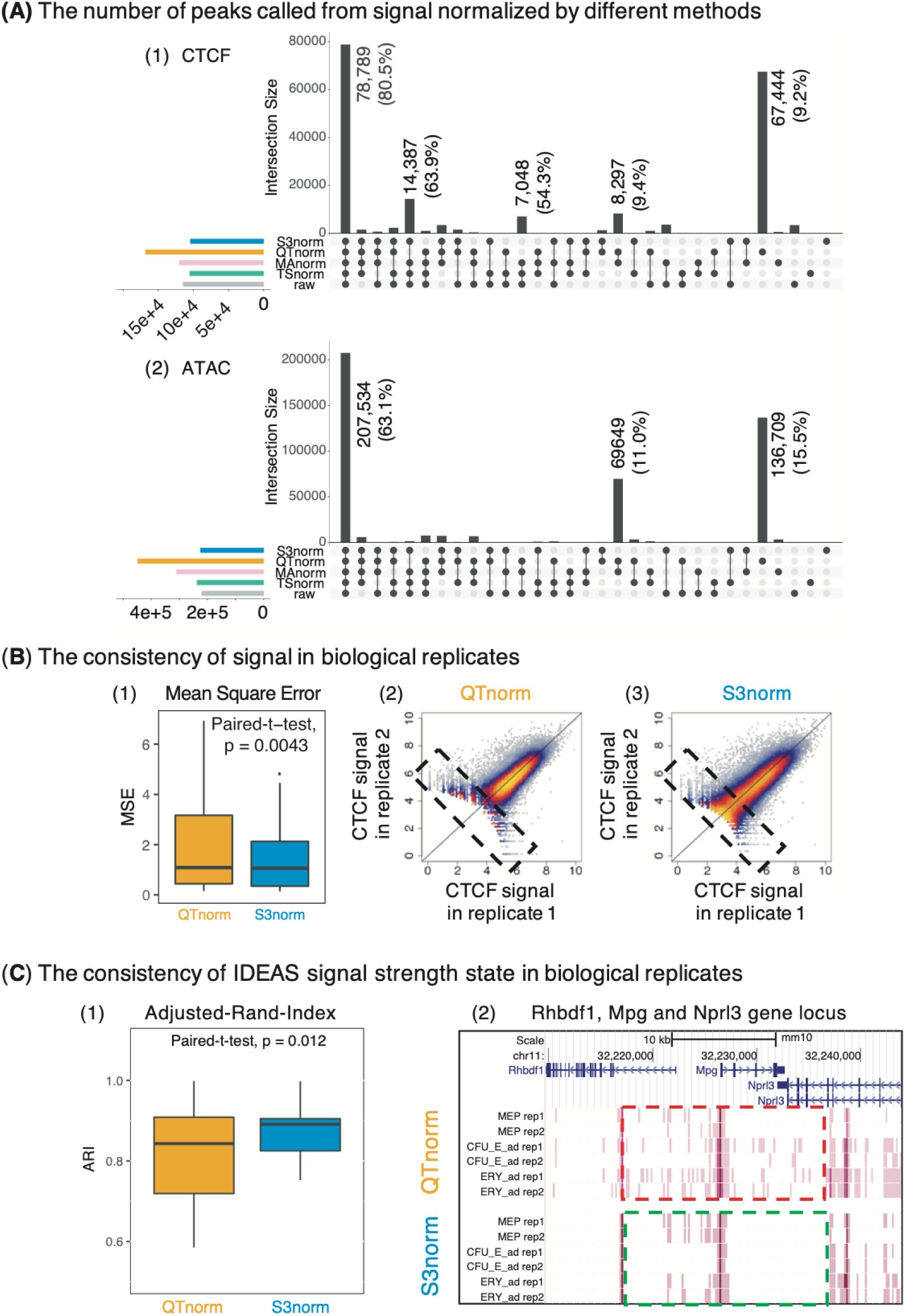
Comparing S3norm and QTnorm by peak calling results, consistencies of signal and signal strength state assignments. **(A)** The UpSet figure of the peak calling results using different normalization signals in VISION project. For each panel, the black bar represents the number of peaks present in the peak calling results by using different normalized signals. The black points below each bar represent the combinations of normalization methods. The left bar represents the total number of peaks called by using a specific normalization method. The (1) panel is the result for CTCF ChIP-seq. The (2) panel is the result for ATAC-seq. For the bars with the substantial number of peaks, the number of the peaks are shown on the top of the bar. For the CTCF ChIP-seq, the percentage of the peaks that include the CTCF motif are shown in the parentheses. For the ATAC-seq, the percentage of the ATAC-seq peaks that intersect with the candidate Cis-Regulatory Elements (cCREs) in the ENCODE SCREEN Project (47) are shown in the parentheses. **(B)** The signal consistency between two biological replicates after normalization by QTnorm and S3norm. Panel (1) shows the boxplot of mean square errors (MSEs) of signals between two biological replicates in all peaks called from the signals normalized by QTnorm and S3norm. Each box includes the MSEs of 15 data sets with biological replicates. The p-value above the boxes represents the significance of paired t-test that tests if the MSEs of S3norm are significantly lower than the MSEs of QTnorm. Two scatterplots (2-3) show the normalized CTCF ChIP-seq signals in two biological replicates. The comparisons of low-signal peaks (counts < 2^6) are highlighted by black dash box. **(C)** The consistency of IDEAS signal strength state assignments between two biological replicates after QTnorm and S3norm. The (1) panel is the boxplot of Adjusted-Rand-Indices (ARIs) of the state assignments between two biological replicates in all ATAC-seq data with replicates. The p-value above the boxes represents the significance of paired t-test that tests if the ARIs of S3norm are significantly higher than the ARIs of QTnorm. The right panel is the screenshot of the IDEAS signal strength states at the Rhbdf1-Mpg-Nprl3 gene locus. The dark pink represents the high signal strength state and light pink represents the low signal strength state. The background state is colored as white. The two dash boxes highlighted the genomic regions showing substantial differences between the assignments of signal strength state in QTnorm and S3norm.

We then used the presence of matches to the CTCF binding site motif to evaluate whether the CTCF peaks found only after normalization by QTnorm were false positives or a reflection of greater sensitivity enabled by that normalization. Given that about 80% of previously determined CTCF binding sites contain the CTCF motif (45, 46), we expected that false positive peaks are less likely to contain a match to the CTCF motif. Indeed, among the QTnorm-specific peaks, only 9.2% had the CTCF binding site motif, whereas 80.5% of the peaks called after normalization by all methods had the CTCF motif. These results suggest that many of the QTnorm-specific CTCF peaks are likely to be false positive peaks.

In a similar evaluation of ATAC-seq datasets, a total of 207,534 peaks were called consistently on the data normalized by all methods (Figure 5A, second panel). Another 136,709 peaks were called only after normalization by QTnorm signal (Figure 5A 2nd panel). We assumed that false positives for ATAC-seq peaks would be less likely to match candidate *Cis*-Regulatory Elements (cCREs) predicted in an independent ENCODE SCREEN project (47). Among the peaks shared by all normalization methods, 63.1% intersected with at least one cCREs in the ENCODE Registry, whereas only 15.5% of the peaks uniquely called after QTnorm normalization overlapped with an ENCODE cCRE. These comparisons suggest that many of the ATAC-seq peaks called on data normalized by QTnorm are likely to be false positive peaks.

We next compared the consistency of signal strengths between biological replicates in peaks called after QTnorm and S3norm normalization, under the assumption that more appropriate normalization will generate greater similarity between replicates. For each replicated CTCF ChIP-seq data set, we pooled and merged the peaks called after QTnorm and S3norm normalization into one union peak list and then aggregated the normalized ChIP-seq signal in each peak in each replicate. Overall, the signals normalized by S3norm showed significantly greater similarity, as shown by the lower values in the distribution of MSE (Figure 5B, first panel). This greater similarity of replicates is accentuated in scatterplots showing much more variance between replicates normalized by QTnorm, especially for the peaks with weaker signals (Figure 5B, second and third panel).

For QTnorm signals, we expect to see lower MSEs in the top peaks because QTnorm assigns the same value to the peaks with the same ranks. This assignment of the same value insures very low MSEs as long as the peaks between replicates have the same rank. Indeed, the QTnorm signals become much less consistent as more peaks were included (Supplementary Figure 5). While normalization by QTnorm gave lower MSEs for the top 30% of peaks, normalization by S3 norm produced significantly lower MSEs, and thus greater replicate consistency, for lists containing the top 50% to 100% of the peaks. Therefore, unless the downstream analysis only focuses on the very top peaks, the S3norm still outperforms QTnorm in terms of signal consistency between biological replicates.

One example of downstream analysis that can be influenced by whole genome data normalization is genome segmentation. This analysis assigns each genomic location to one of a set of discrete states that represent commonly occurring signal strengths and combinations of features, often epigenetic marks. If a normalization procedure introduced a systematic bias between data sets used as input, e.g. by inflating the background, then the segmentation could assign incorrect states in epigenomes. Thus, we compared the impact of normalization by QTnorm and S3norm on epigenetic state assignments. We chose the IDEAS genome segmentation system (19) because it considers a range of signal strengths, not a binary presence or absence call, in state assignments. For this evaluation, we used IDEAS in a simple task of identifying and assigning states determined by the strength of ATAC-seq signals rather than finding states that are combinations of epigenetic features. We assumed that the assignment of consistent ATAC-seq signal strength states between replicates would indicate a more accurate normalization. The ATAC-seq data normalized by S3norm gave significantly more consistent state assignments, as measured by the Adjusted Rand Index (ARI) (Figure 5C panel (1)). This greater replicate similarity is illustrated in the *Rhbdf1*-*Mpg*-*Nprl3* gene locus (Figure 5C panel (2)). While the high signal states are consistent between replicates for both methods, as expected, many inconsistencies between the two biological replicates were observed for the genomic locations assign to the low signal state after normalization by QTnorm. In contrast, the same genomic locations were often assigned to the background state (colored white) after S3norm normalization. These comparisons confirmed that normalization by QTnorm can inflate low signals that are actually background, thereby leading to erroneous state assignment.

These several evaluations of peaks and signal strengths after different methods of normalization show that the S3norm method is advantageous over QTnorm when a broad range of data values are considered, reducing the numbers of false positive peaks and limiting incorrect signal strength state assignments in lower signal regions.

### The influence of hyperparameters on S3norm

We evaluated whether S3norm was robust to the choice of values for its two major hyperparameters. One hyperparameter is the FDR threshold that determines the common peak regions and the common background regions from which S3norm to learns the scale factors in the nonlinear transform, α and β. While the FDR threshold affected the number of the common peak regions and common background regions, it had no significant impact on the values of α and β (Supplementary Figure 6). Another major hyperparameter in S3norm is the bin size of the input data. The epigenomic sequencing data typically have many regions with no signals, and thus, differences in bin size can influence the signal distribution of the input data. To evaluate how robust S3norm is to differences in bin sizes, we determined the values for the scale factors using input data divided into bins of different sizes. We found that the values for β in the S3norm transformation model changed very little with bin size (Supplementary Figure 7A). The scale factor α also showed little differences among bin sizes of 50bp, 100bp, and 200bp, but it did shift to somewhat higher values for particular datasets in 500bp bins. Further analysis revealed that the distribution of signals in 500bp bins was substantially different from the distributions of signals bins of 200 or fewer bp (Supplementary Figure 7B). In summary, these comparisons indicate that S3norm is robust to most changes in either of the two major hyperparameters.

## DISCUSSION

We introduce a simple and robust method to normalize the signals across multiple epigenomic data sets. The essence of this method is to use a nonlinear transformation to rotate the signal of the target data set to that of the reference data set, so that the mean signals of both common peak regions and common background regions are matched simultaneously between the two data sets. The S3norm method achieves several notable improvements over existing normalization methods. First, the inclusion of background regions is a particular advantage when data across the entire genome needs to be normalized. As an example, this method was developed to facilitate our work on genomic segmentations that assign every genomic interval to an epigenetic state, which is a common combination of epigenetic features (20). An inflation of background noise could result in assigning regions with increased noise to low signal-containing states. Second, in contrast to the TSnorm and QTnorm methods, S3norm is robust to biases resulting from the substantial proportion of background regions in the genome. Third, S3norm can be trained on data sets with small numbers of peaks, such as data sets that include spike-in controls (48). Finally, S3norm has only two parameters to be trained from the data, which makes the method robust across a wide variety of data sets.

A key assumption of the S3norm method is that true biological signals should have the same means in common peak regions and in common background regions between data sets. For some data sets in which different signal strengths in common peak regions are expected, a variation on the S3norm method may be more appropriate. For example, for ChIP-seq data sets of transcription factors whose abundance is changing over the course of a targeted degradation protocol, we expect the mean signals in peak regions (which are common peak regions across the time course) to deteriorate over time. In such cases, one should not use any of the data sets in the time course as a reference for S3norm normalization. Instead, a spike-in control or a small number of unchanged peak regions identified by other techniques should be paired with the background regions at each time point in order for S3norm to work properly.

In summary, S3norm is a simple and robust method to normalize multiple data sets. The results of applying S3norm to epigenomic data sets show that it is more effective in bringing out real biological differences than existing methods. As more epigenomic data continue to be generated, S3norm will be useful to normalize signals across these diverse and heterogeneous epigenomic data sets to allow downstream analyses to capture true epigenetic changes rather than technical bias. Improved normalization will aide studies that analyze data sets across multiple experiments, such as differential gene regulation, genome segmentation (17, 19), joint peak calling (49), predicting gene expression (50), and detecting transcription factor binding events (51).

## Supporting information

S3norm_manuscript_supplementary_Figures_and_Methods_revision

## DATA AVAILABILITY

Files for raw signals, p-value converted signals, and signals from S3norm are available both for download and for viewing from the VISION website (http://usevision.org). The S3norm normalization package is available at GitHub (https://github.com/guanjue/S3norm).

## ACKNOWLEDGEMENT

We thank Dr. Alexander Q. Wixom and Tiffany Wang for their help to test S3norm package.

## FUNDING

The work is supported by National Institutes of Health grants [GM121613 to Y.Z.], [DK106766 to Y.Z. R.C.H., and Q.L.], [GM109453 to Q.L.] and [CA178393 to R.C.H.].

