## Supplementary material for "S3norm: simultaneous normalization of sequencing depth and signal-to-noise ratio in epigenomic data": S3norm_manuscript_supplementary_Figures_and_Methods_revision

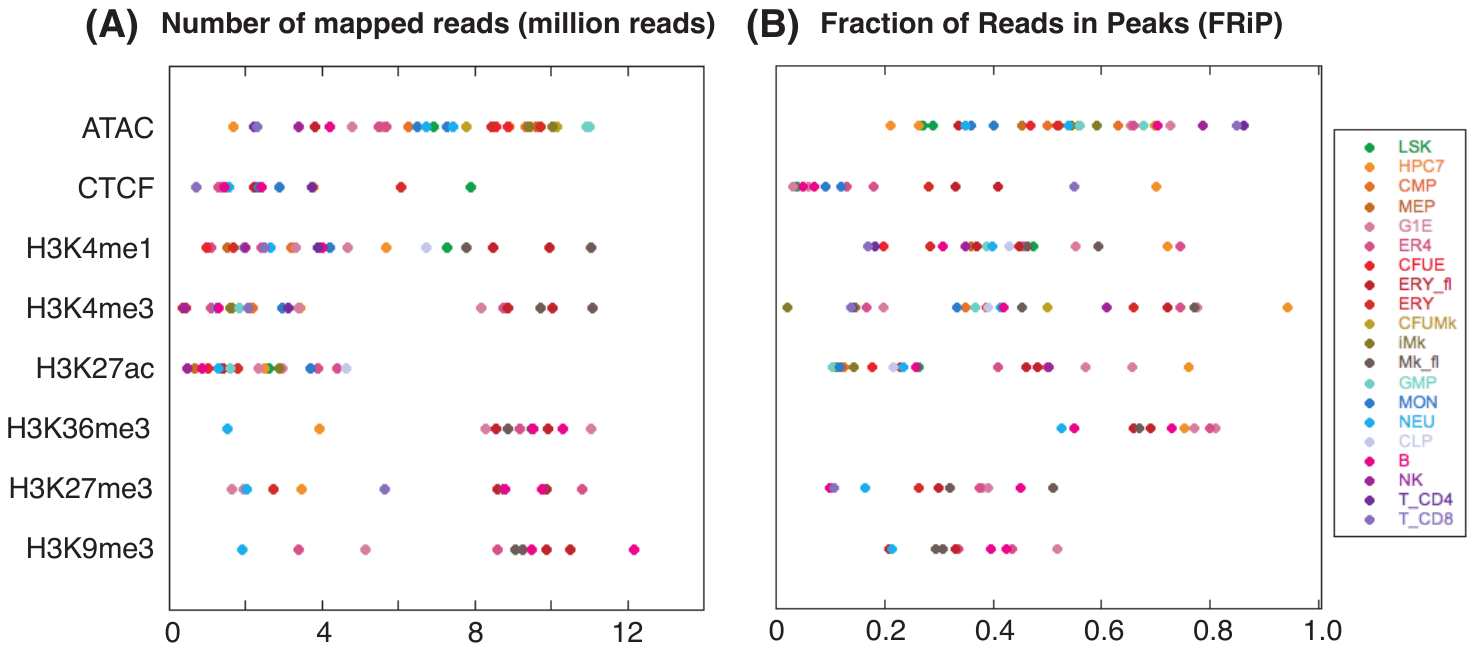

**Supplementary Figure 1.** Demonstration of data heterogeneity in the **V**al**I**dated and **S**ystematic integrat**ION** of epigenomic data project (**VISION**: usevision.org). There are to eight epigenetic marks in twenty hematopoietic cell types in the VISION project. **(A)** The number of mapped reads of each data set. **(B)** The FRiP score of each data set. Each point represents one data set. The color of the point represents the cell type of the data set.

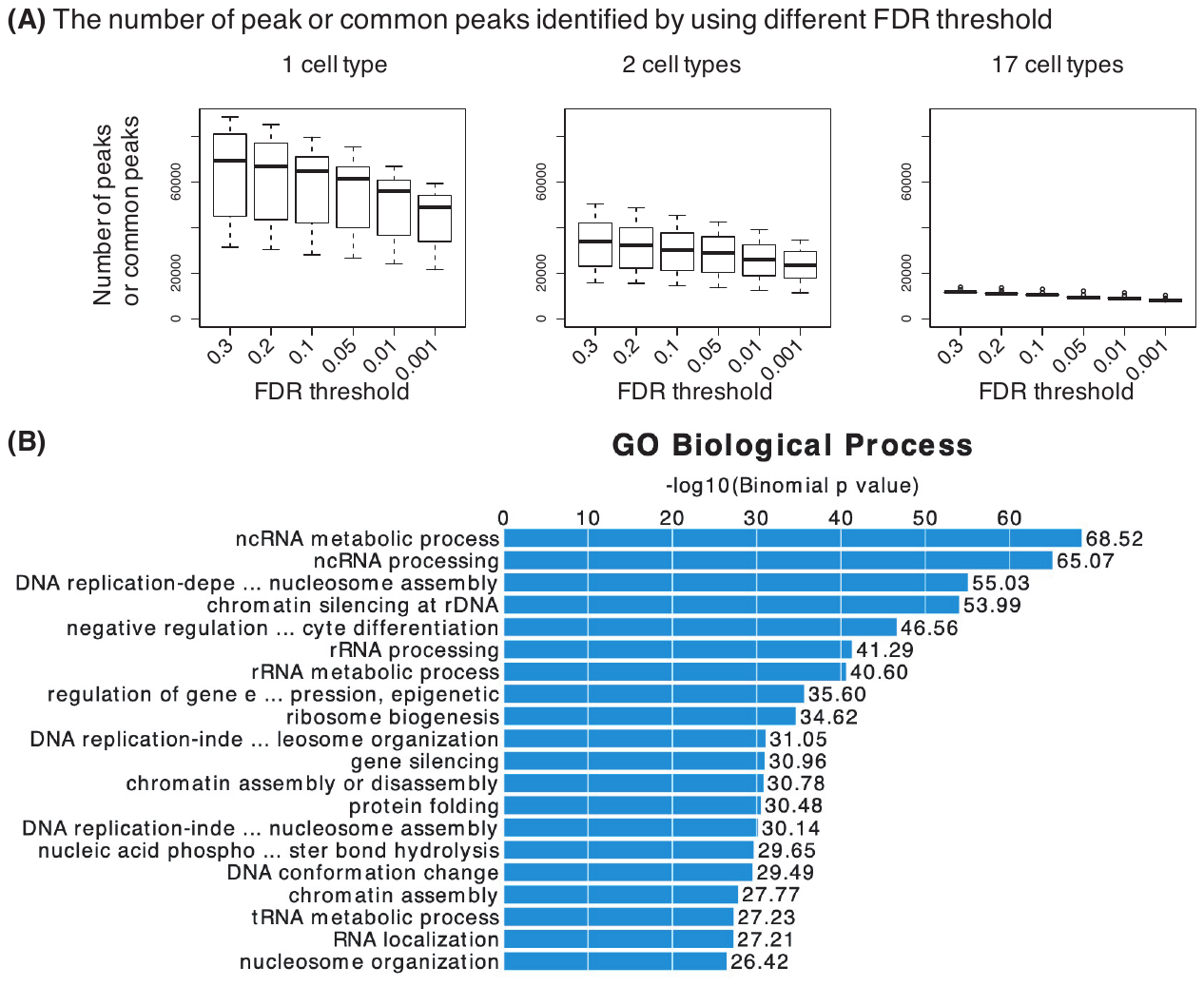

**Supplementary Figure 2.** (A) The numbers of ATAC-seq peaks called by using different FDR thresholds. The first panel is showing the boxplot of peaks called in each cell type. The second panel and third is showing the number of common peaks shared by 2 cell types and 17 cell types. (B) The gene ontology associated with ATAC-seq common peak regions (FDR < 0.01) shared by 18 cell types in VISION project.

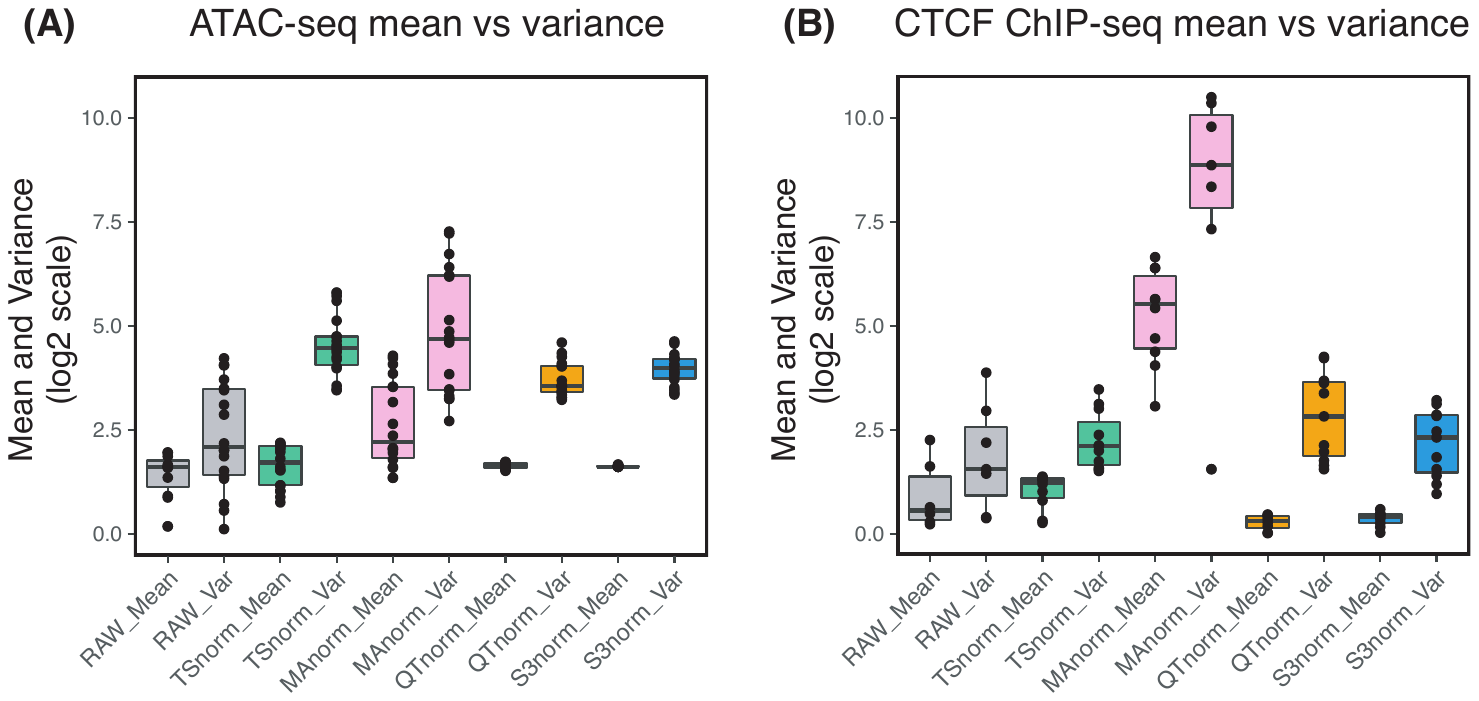

**Supplementary Figure 3.** Evaluation of variance inflation in the background signal in **(A)** 18 nuclease sensitivity (ATAC-seq or DNase-seq) data sets and **(B)** 11 CTCF ChIP-seq data sets. The different normalization methods are arrayed along rhe x-axis, and the y-axis displays the mean and the variance of the data sets (log2 scale). For each method, the first box is the mean of the background signal and the second box is the variance of the background signal.

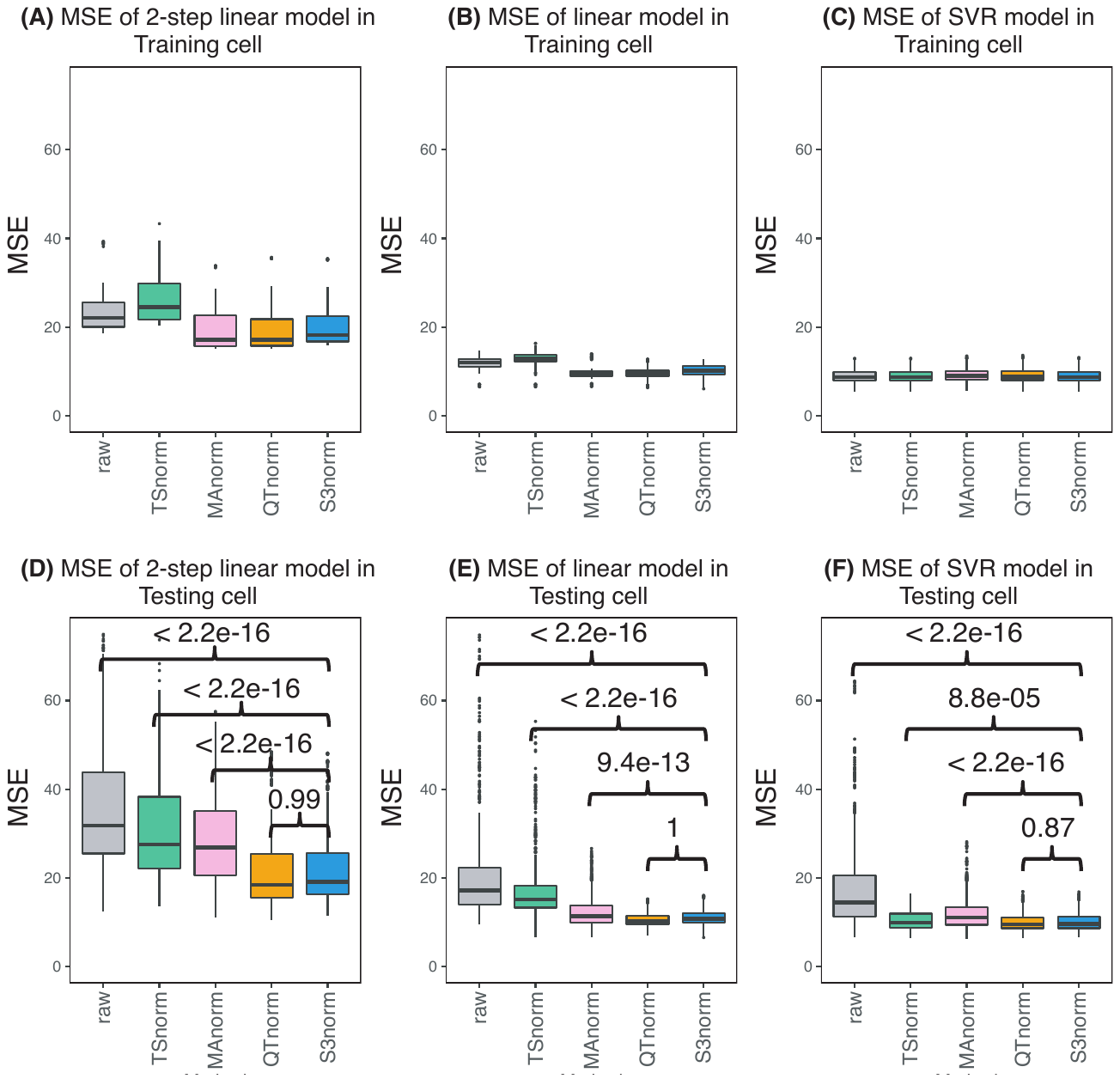

**Supplementary Figure 4.** Comparison of S3norm and other methods by the ability of normalized histone modification data to predict gene expression. (**A-C**) The MSE of the observed RNA-seq signal and the predicted RNA-seq in ten-fold cross validation in the **Training Cell Type** by using 2-step linear regression model (**A**), linear regression model (**B**), and SVR model (**C**). (**D-F**) The MSE of the observed RNA-seq signal and the predicted RNA-seq in ten-fold cross validation in the **Testing Cell Type** by using 2-step linear regression model (**D**), linear regression model (**E**), and SVR model (**F**). The p-values above the boxes come from the Wilcoxon test that tests if the MSE of S3norm are significantly better than the other methods.

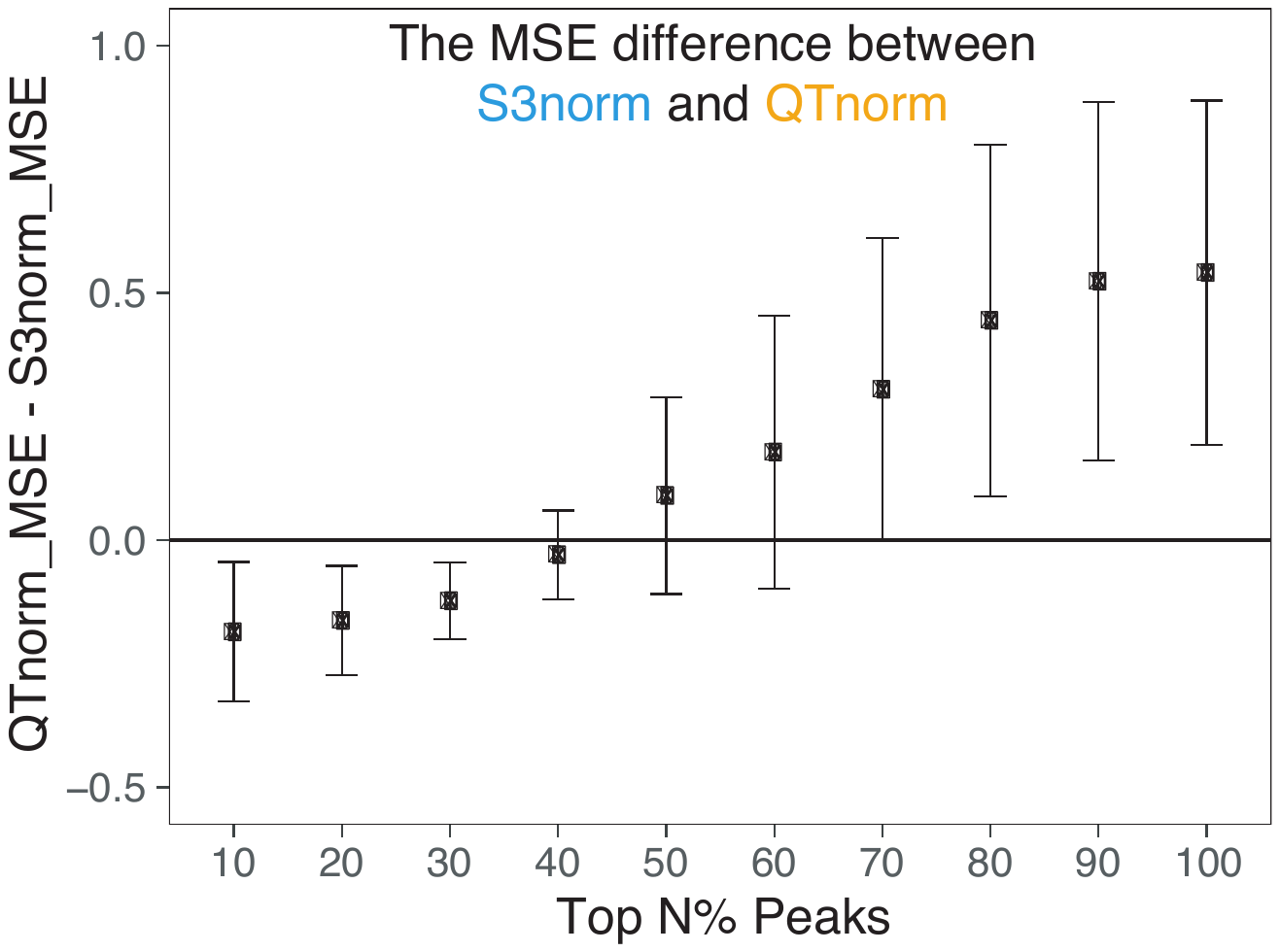

**Supplementary Figure 5.** The difference of MSEs between signals in biological replicates after normalization by QTnorm and S3norm. Each point represents the average difference of MSE in 15 datasets with biological replicates. The error bar represents the 95% confidence interval of the differences of MSE.

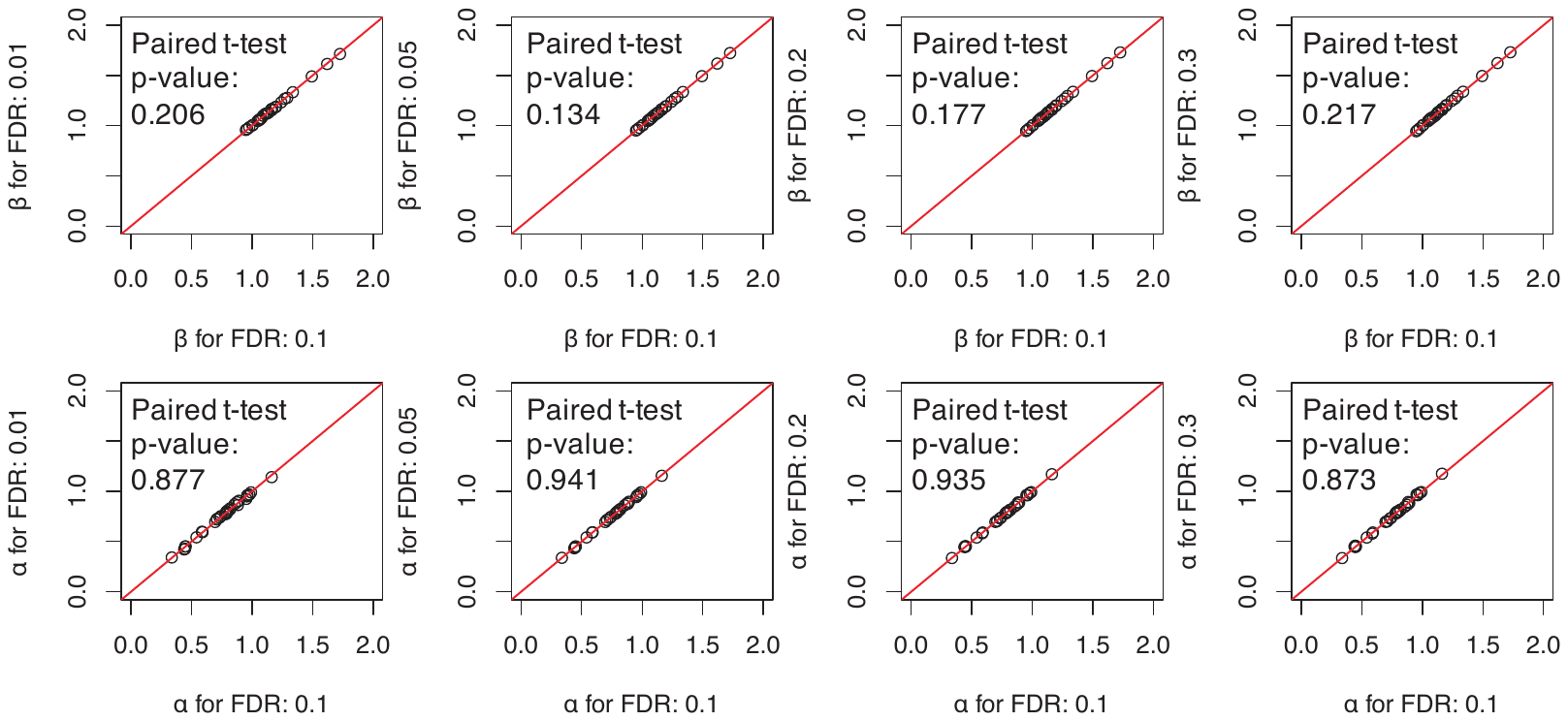

**Supplementary Figure 6.** The robustness of S3norm to differences in FDR threshold. The two scale factors of S3norm, α and β, were learned from common peak regions and common background regions, which were determined at different FDR thresholds. The values of the scale factors were evaluated in a series of pairwise comparisons, in which the x-axis shows the scale factors learned from common peak regions and common background regions determined by 0.1 FDR, and the y-axis shows the scale factors learned from the common peak regions and common background regions determined by other FDR thresholds (0.01, 0.05, 0.2, 0.3). The red line is y=x line. The first and second rows of graphs show the comparisons for the β’s and α’s, respectively.

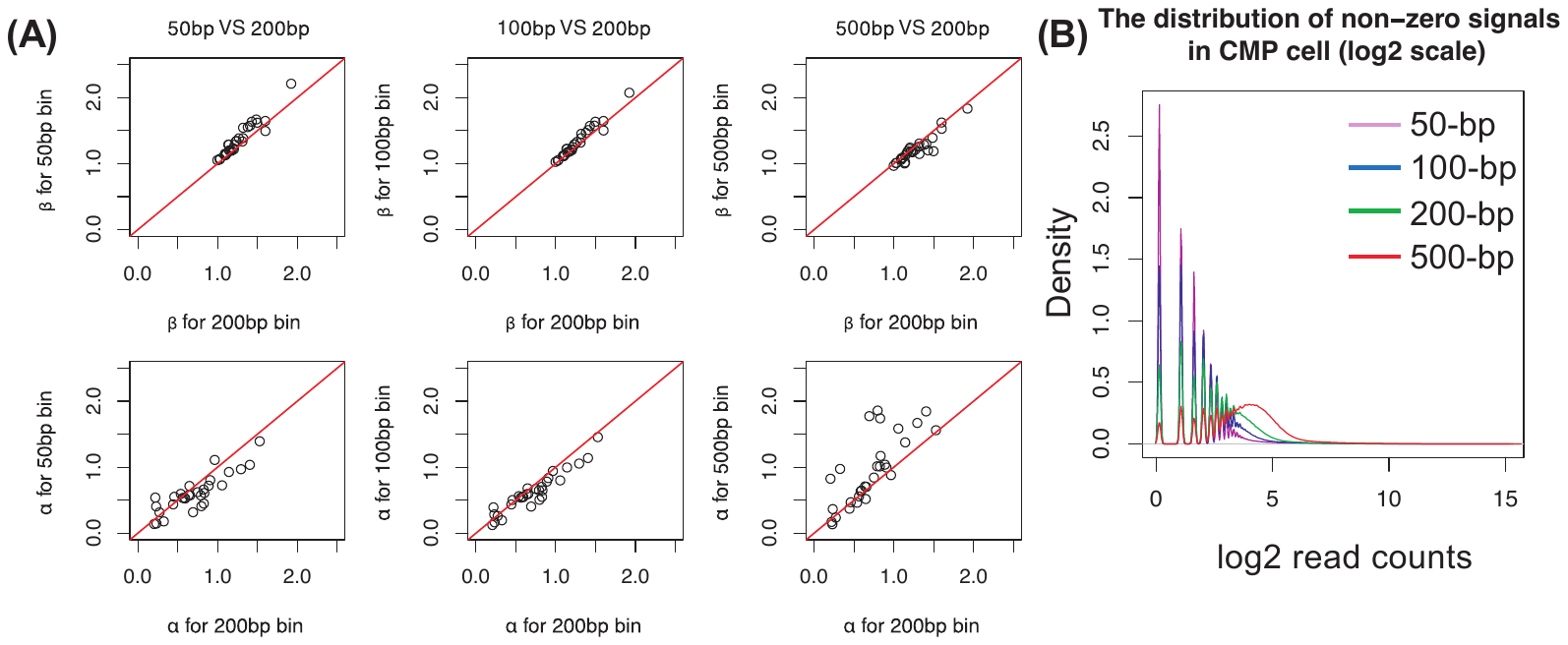

**Supplementary Figure 7.** (**A**) The robustness of S3norm relative to the bin size of the input data. In each graph, the x-axis shows the scale factors β (top row) or α (bottom row) learned from the signal in 200bp bins. The y-axis shows the scale factors learned from signal based on other bins sizes (50bp, 100bp, 500bp). The red line is y=x line. (**B**) The distribution of non-zero read counts in epigenetic datasets from the CMP cell population. Each line represents the distribution of non-zero read counts in bins of the designated size.

**SUPPLEMENTARY METHODS**

### **Estimating NB model from non-zero data**

For epigenomic sequencing data, there are often large number of zero bins across the whole genome (1). To avoid potential problem caused by this issue, we estimate the probability of success parameter (p) and the shape parameter ($s_{\mathrm{local}}$) in the dynamic NB model by only using the non-zero data.

Let N denote the number of non-zero bins, p0 and N0 denote the proportion of zero bins across the whole genome, M1 and M2 denote the mean of the data and mean of square of the data, NM1 and NM2 denote the non-zero mean and non-zero mean of square of the data.

$M1=\frac{NM1\times N+0\times N0}{N+N0}$	(1)

$M1=NM1\times\frac{N}{N+N0}$	(2)

$M1=NM1\times\left( 1-p0 \right)$ (3)

$M2=\frac{NM2\times N+0\times N0}{N+N0}$	(4)

$M2=NM2\times\frac{N}{N+N0}$	(5)

$M2=NM2\times\left( 1-p0 \right)$ (6)

The variance can be calculated by subtracting the square of the mean of the data from the mean of square of the data (2).

$\sigma^{2}=M2-{M1}^{2}$	(7)

Then, the p and the $s_{\mathrm{local}}$ in the dynamic NB model can be defined as follows:

$p=\frac{\sigma^{2}-M}{\sigma^{2}}$	(8)

$p=1-\frac{M1}{M2-{M1}^{2}}$	(9)

$p=1-\frac{NM1\times\left( 1-p0 \right)}{NM2\times\left( 1-p0 \right)-{{NM1}^{2}\times\left( 1-p0 \right)}^{2}}$	(10)

$p=1-\frac{NM1}{NM2-{NM1}^{2}\times\left( 1-p0 \right)}$ (11)

$s_{\mathrm{local}}=\frac{{M1}^{2}}{\sigma^{2}-M}\times\frac{r_{i}^{\mathrm{ctrl}}}{M^{\mathrm{ctrl}}}$	(12)

$s_{\mathrm{local}}=M1\times\frac{1-p}{p}\times\frac{r_{i}^{\mathrm{ctrl}}}{M^{\mathrm{ctrl}}}$ (13)

$s_{\mathrm{local}}=NM1\times\left( 1-p0 \right)\times\frac{1-p}{p}\times\frac{r_{i}^{\mathrm{ctrl}}}{M^{\mathrm{ctrl}}}$ (14)

Since expected p0 without zero-inflation can be calculated by the p and $s_{\mathrm{local}}$, those parameters can be estimated by iteratively updating the three parameters by using the three equations:

$p0=\left( 1-p \right)^{s}$ (15)

$p=1-\frac{NM1}{NM2-{NM1}^{2}\times\left( 1-p0 \right)}$ (11)

$s_{\mathrm{local}}=NM1\times\left( 1-p0 \right)\times\frac{1-p}{p}\times\frac{r_{i}^{\mathrm{ctrl}}}{M^{\mathrm{ctrl}}}$ (14)

The p0 is initialized as the observe proportion of zero bins in the data.
